# LAB-AID (Laboratory Automated Interrogation of Data): an interactive web application for visualization of multi-level data from biological experiments

**DOI:** 10.1101/763318

**Authors:** Zrinko Kozic, Sam Booker, Owen Dando, Giles Hardingham, Peter Kind

**Affiliations:** Simons Initiative for the Developing Brain, University of Edinburgh, Edinburgh, United Kingdom; Patrick Wild Centre, University of Edinburgh, Edinburgh, United Kingdom; UK Dementia Research Institute, University of Edinburgh, Edinburgh, United Kingdom; Centre for Discovery Brain Sciences, University of Edinburgh, Edinburgh, United Kingdom

## Abstract

A key step in understanding the results of biological experiments is visualization of the data. Many laboratory experiments contain a range of measurements that exist within a hierarchy of interdependence. An automated way to visualise and interrogate experimental data would: 1) lead to improved understanding of the results, 2) help to determine which statistical tests should be performed, and 3) easily identify outliers and sources of batch effects. Unfortunately, existing graphing solutions often demand expertise in programming, require considerable effort to import and examine such multi-level data, or are unnecessarily complex for the task at hand. Here we present LAB-AID (**Lab**oratory **A**utomated **I**nterrogation of **D**ata), an interactive tool specifically designed to automatically visualize and query hierarchical data resulting from biological experiments.

## Introduction

There is a great demand for biologists to be able to visualize data from their laboratory experiments. This is particularly true when measurements exist within a hierarchy of interdependence, and when multiple sources of batch effects may influence experimental results. For example, in an investigation into the electrophysiological properties of neurons in the brain of a transgenic rodent model of a particular neurodevelopmental or neurodegenerative condition, diverse measurements are made on individual neurons whose behaviour is not independent of each other (e.g. they were subject to the same developmental program that gave rise to a functioning brain). From a single animal, multiple neurons may be probed within an individual brain slice or multiple slices. Furthermore, multiple animals will be tested which may or may not be litter mates or have shared cages. Random effects (such as slice, animal or litter) at any level within this hierarchy can consistently impact groups of measurements leading to misinterpretation of the results. Moreover, data may be gathered by different investigators on several dates using diverse experimental equipment, resulting in potential batch effects that may further complicate interpretation. These random and batch effects can be difficult to disentangle from the factor being investigated. In addition, it is often important to understand the relationship or correlation between different measurements made on the same experimental unit (for example, multiple different electrophysiological measurements made on each neuron, or multiple behavioural measurements made on each animal).

Thus, it is important to thoroughly visualize and understand such data before proceeding to statistical tests of effects, otherwise misleading, inappropriate or even spurious results may be obtained. The proper visualisation of the data can also help researchers guard against common statistical errors (e.g. pseudoreplication). In addition, visualization allows for the identification of outliers at all levels within the hierarchy of measurements, and may help to determine cases of erroneous measurement or recording of data.

While numerical and statistical computing environments such as R [1] and MATLAB [2] provide a rich and powerful framework within which experimental data can be examined and plotted, their use is not necessarily straightforward for those without experience or training in programming. Dedicated scientific graphing software tools such as GraphPad Prism [3] provide graphical user interfaces which may be more approachable for the laboratory scientist, a number of online tools have recently been developed to automatically plot experimental data provided in the form of a spreadsheet or table (for example [4], [5], [6], [7], [8]), and more general online plotting tools are available (such as Plotly Chart Studio [9] and RAWGraphs [10]). However, such software may be more oriented towards examining a single experimental variable at a time, to producing publication-ready plots rather than querying data, or towards providing a wide-range of general features and plot types at the price of some complexity of use.

Hence it would be of considerable use to laboratory scientists to have a tool which could automatically plot such multi-level data from biological experiments, by which they could examine distributions, correlations between variables, outliers and batch effects. Here we present LAB-AID (**Lab**oratory **A**utomated **I**nterrogation of **D**ata), a tool specifically designed to visualize and query multi-level data resulting from biological experiments, that is easy to install, configure and use.

## Results

As a demonstration of LAB-AID, we use experimental data in which the morphology, function and integration of dendritic spines was studied in layer 4 spiny-stellate cells from the somatosensory cortex of a mouse model of a monogenic neurodevelopmental disorder, and compared with the behaviour of neurons of wild-type (control) mice [11]. In the particular data shown, intrinsic electrophysiological properties were measured on neurons from control and knockout mice; multiple cells were tested per brain slice, and multiple brain slices per experimental animal. Data sets for LAB-AID take the form of an Excel spreadsheet or comma-separated values (CSV) file, in which each row comprises metadata and measurements of experimental variables on a single experimental unit (for example, a cell); multiple such data sets may be loaded into the application at the same time.

The LAB-AID user-interface is split into seven major sections (see Fig 1), available from the main tab bar, with different functions:

1. **Data Select**: Choose, depending on their metadata, which data points will be plotted, and which highlighted. Preview data selections and highlighting.
2. **Data Plots**: View box or violin plots, bar plots, histograms or density plots, or cumulative frequency distributions of chosen data, comparing between and across levels of fixed and random effects.
3. **Modelling**: Perform simple linear (mixed) model fitting and test the significance of fixed effects.
4. **Correlation**: View correlations between experimental variables, calculated across experimental units, and comparisons of correlations between different levels of particular effects.
5. **Downloads**: Export subsets of the input data for download.
6. **Configuration**: Configure the data sets available for investigation, and modify the application title and description.
7. **About**: Read information describing the presented data sets.

**Fig 1.**
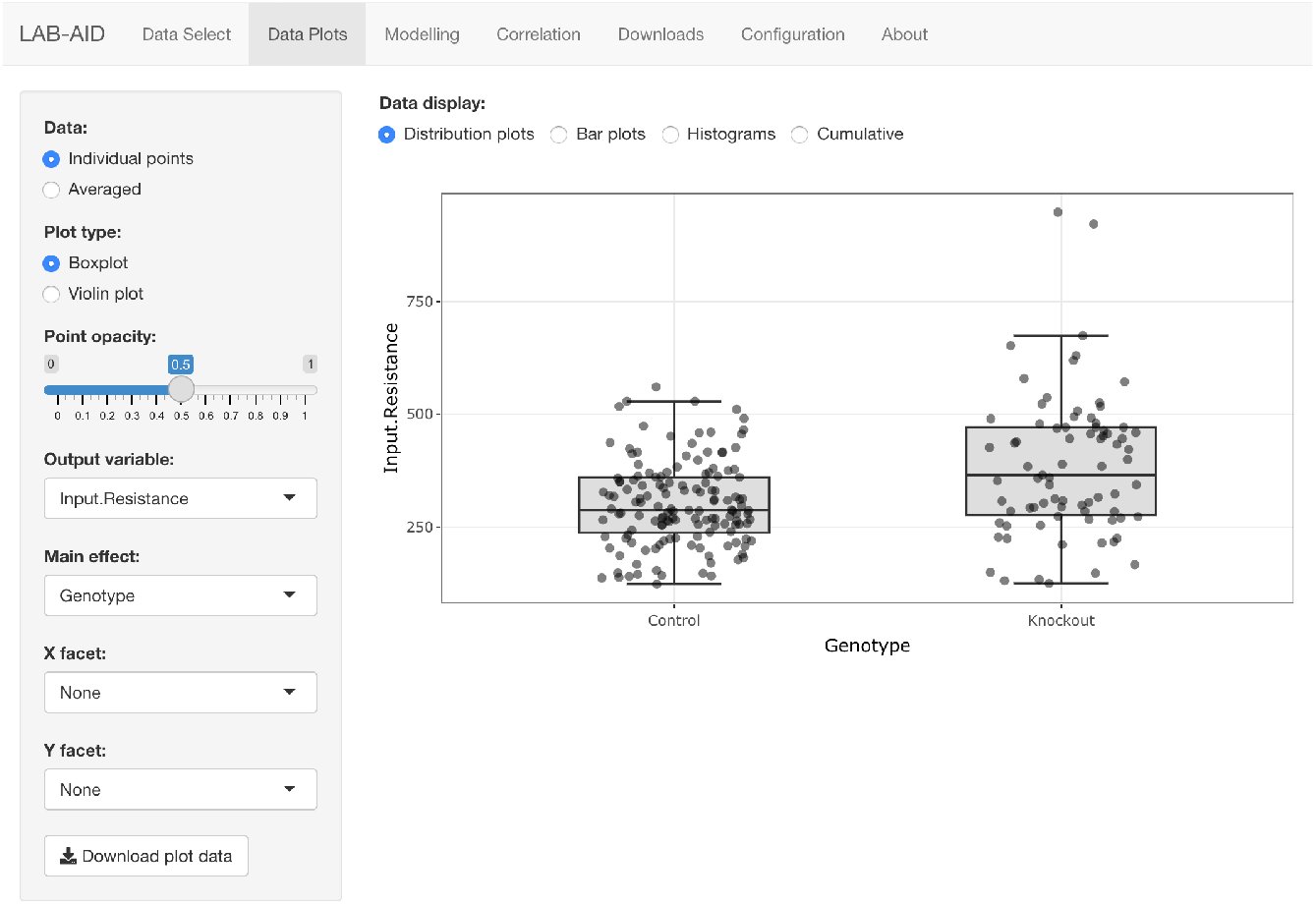
The LAB-AID interface is split into seven major sections. The **Data Select**, **Data Plots**, **Modelling**, **Correlation**, **Downloads**, **Configuration** and **About** sections are accessible from the main tab bar. Here the **Data Plots** tab is selected.

### Data selection and highlighting

A drop-down selection box at the top of the **Data Select** tab allows the user to choose the data set (of those currently loaded) to be examined. Below this, four further tabs give access to data selection, highlighting, preview and summary statistics functionality.

Next to the **Dataset** selection box is a multiple selection drop-down box, **Averaging factors**, which allows the user to select metadata factors over which the measured variables can be averaged. A common use case would be to allow measurements to be shown averaged for individual animals (i.e. animal means), but it also possible to average over different experimental conditions, such as experimental dates, or animal models. To achieve averaging over brain regions within each animal in the example data set, all factors except *slice* and cell should be selected. Whether individual or average measurements will be plotted can be selected in the sidebar menu on the **Data Plots** tab (see below).

#### Selecting data to be plotted

The **Selection** sub-tab allows the user to choose which data are to be included in graphs (see Fig 2a). A drop-down selection box is present for each column of metadata in the current data set. These allow subsets of the data to be included in or excluded from plots, depending on the metadata characteristics of each experimental unit. Individual levels of each metadata factor can be selected or deselected individually; in addition, **Select All** and **Deselect All** buttons allow for bulk selection and deselection. By default, all data in the data set is selected to be shown in plots.

**Fig 2.**
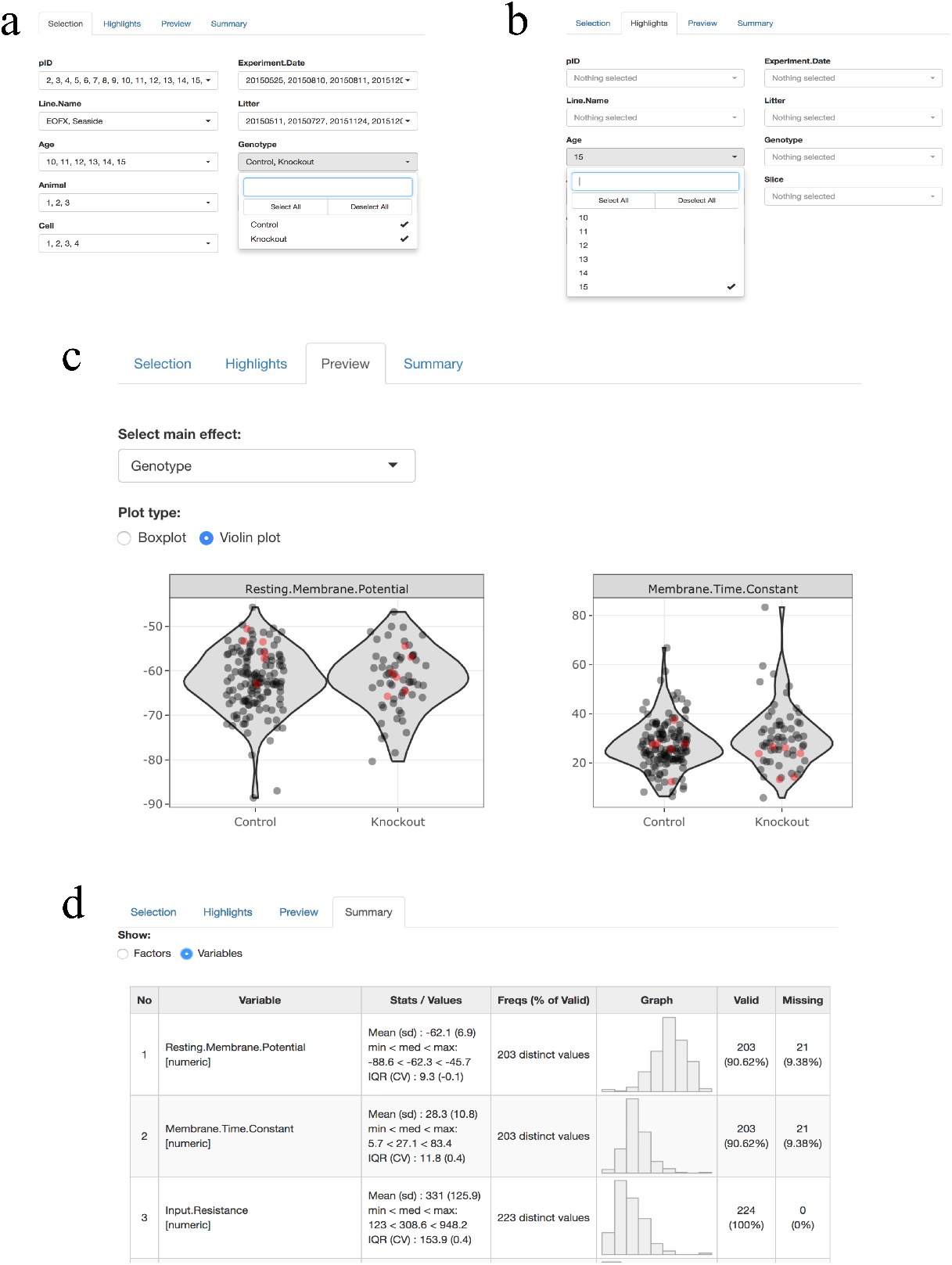
The Data Select tab allows selection of the data to be displayed and highlighted, previews those choices, and displays summary statistics. (a) The Selection sub-tab includes or excludes data subsets from plots, based on the metadata characteristics of each experimental unit. (b) The Highlights sub-tab allows data subsets to be highlighted in colour, based on the metadata characteristics of each experimental unit. Here, data from cells originating from animals aged 15 days are to be highlighted. (c) Selection and highlighting choices can be inspected in the Preview tab, which also presents an overview of the distribution of data for all experimental variables within a particular data set. Here, data from cells originating from animals aged 15 days have been highlighted in red. (d) Summary statistics for the data are shown in the Summary sub-tab. Statistics respect the data selection choices made in the Selection sub-tab.

#### Highlighting data subsets

The **Highlights** sub-tab allows the user to choose which data, among that included in the graphs, is to be further highlighted in colour (see Fig 2b). Its user interface is analogous to that of the **Selection** tab, with a drop-down selection box for each column of metadata in the current data set; data points are highlighted according to the metadata characteristics of each experimental unit. By default, no data are selected to be highlighted.

#### Previewing data plots

The **Preview** sub-tab presents a simple plot of the distribution of data for each experimental variable within the current data set. Individual data points are shown (see Fig 2c), and overall distributions can be represented as either box or violin plots. Data are graphed according to the different levels of a single effect, chosen from the metadata columns in the input data. Preview plots respect the selection and highlighting choices made in the **Selection** and **Highlights** tabs.

Hovering over any data point in the **Preview** tab (or in the main **Data Plots** tab) will display a tooltip showing metadata attributes for that data point. In addition to the metadata derived from the input Excel or CSV file, a point ID (pID) attribute is shown, which corresponds to the relevant row in the input file, thus allowing easy location of points of interest within the input data. The pID attribute can also be used to easily highlight particular points, or exclude them from plots.

#### Displaying summary statistics

The **Summary** sub-tab displays a table of summary statistics for the data (see Fig 2d). If the **Factors** radio button is selected, the table displays a list of all the experimental factors from the data set, along with up to ten levels of each factor, their corresponding number of data points, frequency in the data set and the number of valid (non-missing) values. Selecting the **Values** radio button shows a table containing all measured variables with summary statistics for each. These statistics include mean, standard deviation, median, minimum and maximum values, inter-quartile range and coefficient of variation. A simple histogram of the each variable’s distribution is plotted, together with a summary of valid, missing and distinct values.

### Data plots

The behaviour of particular experimental variables can be examined in detail in the top-level **Data Plots** tab. The main area of the tab shows either (i) data distribution plots, (ii) bar plots, (iii) histogram or density plots, or (iv) cumulative frequency plots, for a particular experimental variable of interest; choice between these four modes is made via the **Data display** radio buttons. At the left of the **Data Plots** tab, a sidebar provides further control over the plots shown in the main area. Some sidebar elements are common to all plot types, whereas others change depending on the plot type.

Among the common controls, the **Data** radio buttons determine whether the data plotted is for individual experimental units (i.e. for each row in the input data file), or if it should be averaged over the levels of metadata factors selected in the **Averaging factors** selection box under the **Data Select** tab. The last four selection boxes control which variable will be plotted (**Output variable**) and how data should be sub-divided for comparison. The **Main effect** selection box determines the metadata factor of primary interest; how this is represented depends on the type of plot being shown, and will be further described below. The **X facet** and **Y facet** selection boxes allow for the further subdivision of the data into separate columns and rows of graphs, respectively, according to the levels of two additional metadata factors.

#### Distributions

If the **Distribution plots** radio button is selected, then individual data points (or averages) for the chosen experimental variable are plotted, along with overall distributions (see Fig 3a); data selection and highlighting choices made in the **Data Select** tab are respected. In each graph displayed, data are separated on the x-axis into separate levels of the main effect chosen in the left sidebar, and the overall distribution shape is shown for each. If no selections are made in the **X facet** and **Y facet** selection boxes, then a single graph is shown containing all data. In the sidebar, radio buttons specific to this plot type control whether overall distributions will be represented as box or violin plots.

**Fig 3.**
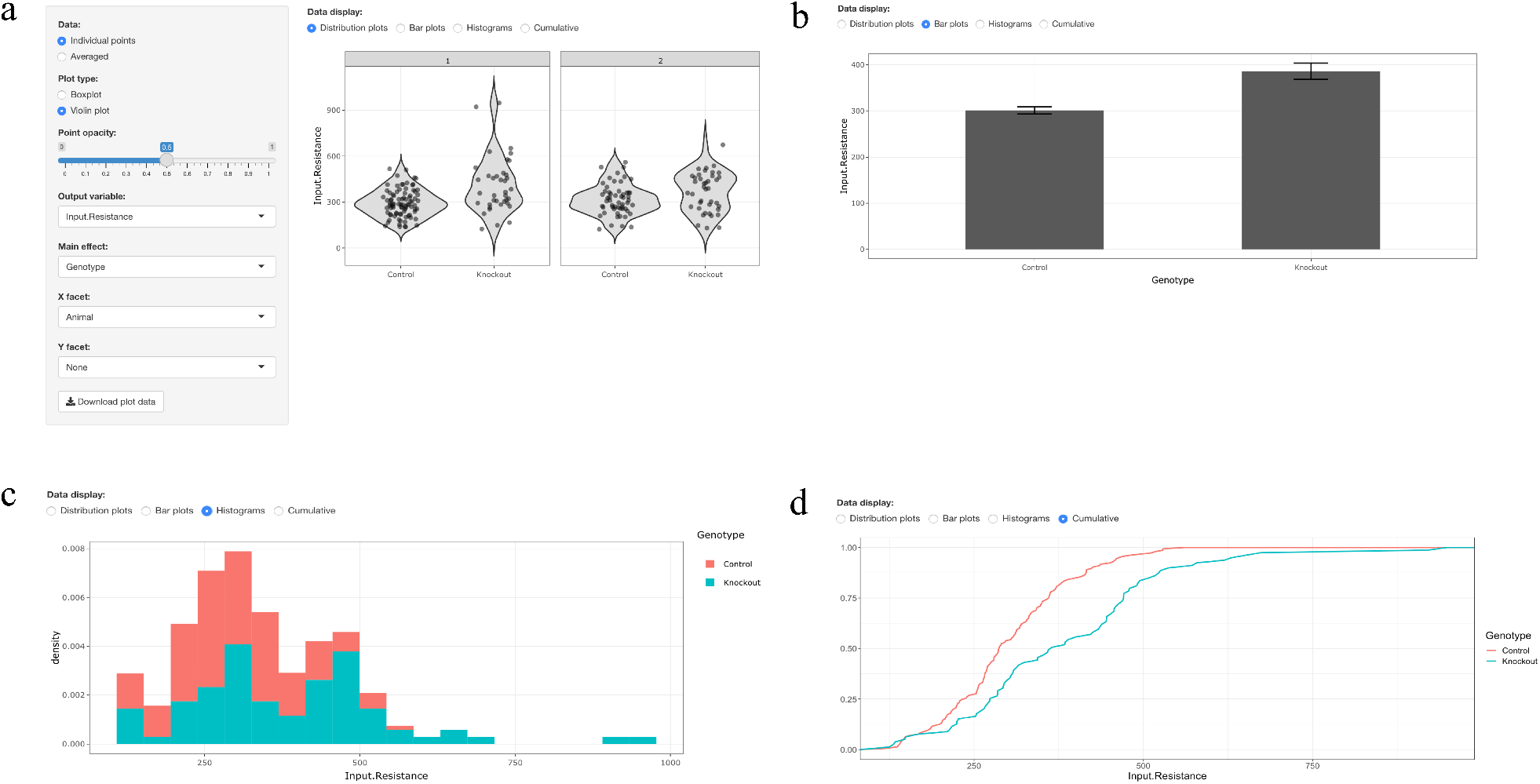
The behaviour of experimental variables can be examined in detail in the Data Plots tab. (a) Distribution plots show individual data points together with box or violin plots for a particular experimental variable. In our example data set, neuronal input resistance is plotted across the two levels of the Genotype effect (i.e. for control or knockout mice). Animal number has been chosen as the X facet, and hence the data are subdivided into a row of separate graphs for each value of this factor. (b) Representation of the mean and SEM for neuronal input resistance as a bar plot. (c) Data can also be represented as histogram or density plots. Here, a histogram of neuronal input resistance is plotted for the two levels of the Genotype effect. (c) Representation of data as a cumulative frequency distribution. In our example data set, neuronal input resistance is plotted for the two levels of the Genotype effect.

#### Bar plots

Selecting the **Bar plots** radio button displays summary statistics for the selected experimental variable in a form of a bar plot, where the height of the bar corresponds to the mean value of the variable. Error bars can be set to display either the standard error of the mean (SEM) or a 95% confidence interval (CI) by selecting the corresponding option in the sidebar menu. Similarly to distribution plots, bars are initially split by the main effect and the plot can be faceted by additional variables. Bar plots can also be generated based on individual points, or averaged over factors selected in the **Averaging factors** selection box.

#### Histograms

When the **Histograms** radio button is selected, data for the chosen experimental variable are plotted as a histogram or density plot, as determined by the **Plot type** radio buttons in the sidebar (see Fig 3b). If data are plotted as a histogram, then the **Bins** slide control in the sidebar determines the number of bins. Data are separated by colour of bars (histogram) or line (density plot) into different levels of the main effect chosen in the sidebar. If no selections are made in the **X facet** and **Y facet** selection boxes, then a single histogram or density plot is shown representing all data, otherwise the data can be subdivided into multiple histograms or density plots according to the levels of one or two extra factors.

#### Cumulative frequency distributions

Finally, if the **Cumulative** radio button is selected, data for the chosen experimental variable are plotted as a cumulative frequency distribution (see Fig 3c). Data are separated by colour of line into different levels of the main effect chosen in the sidebar. As before, a single plot encompassing all data can be shown, or the data can be subdivided according to one or two additional factors. There are no plot-type specific controls in the sidebar for cumulative frequency distributions.

### Modelling

LAB-AID allows users to perform simple statistical testing on data sets by fitting linear (mixed) models. While this functionality should not replace detailed modelling and analysis, which requires statistical expertise, it can allow users to quickly assess the likely magnitude of effect sizes, and to investigate the influence of modelling the levels of the hierarchy of interdependence of their data on calculations of statistical significance.

In the top-level **Modelling** tab, linear model and linear mixed model fitting are performed, respectively, using the lm and lmer functions from the lme4 R package [12]. Which variable is to be modelled can be chosen from **Select variable** drop-down menu (see Fig 4a). Selecting the **Log-transformed** checkbox will log-transform this variable (and if the data contains any non-positive values, a warning will be displayed to indicate that the log-transformation cannot be applied). Below these data selection and transformation options a quantile-quantile (QQ) plot of the selected variable is displayed. The QQ plot can help to assess how well the raw or log-transformed data fits a normal distribution; data with error distributions other than the normal distribution would require fitting to a generalized linear or generalized linear mixed model, which is beyond the scope of LAB-AID’s functionality.

**Fig 4.**
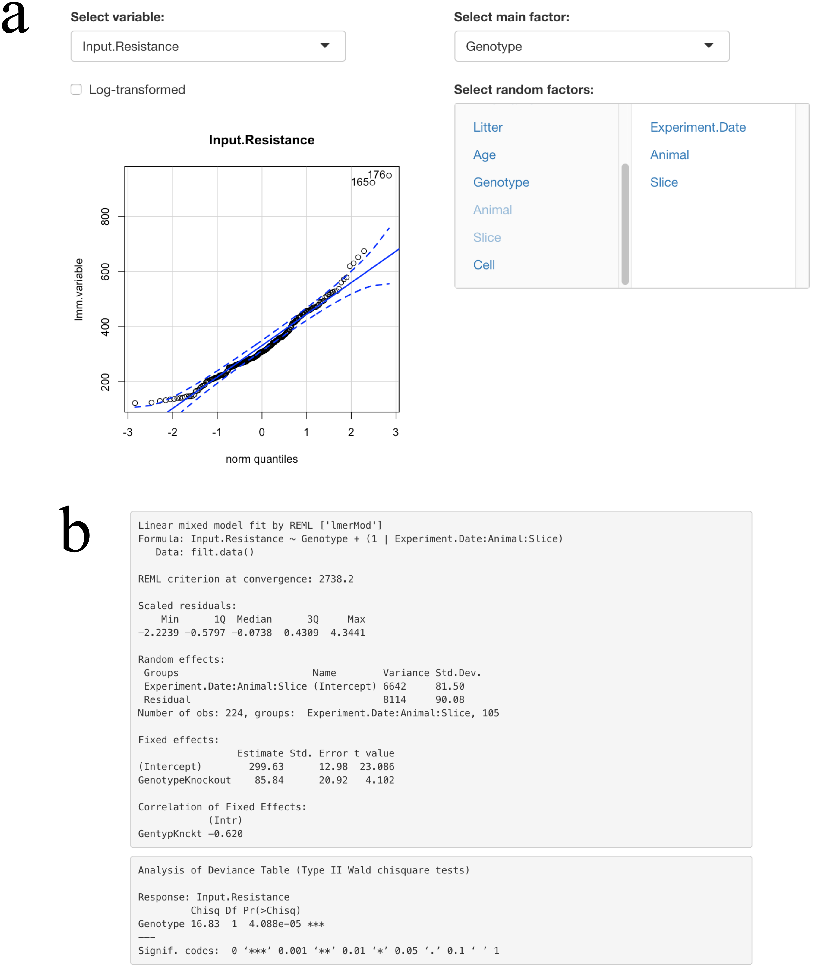
The influence of fixed and random effects on a variable can be assessed in the Modelling tab. (a) Drop-down lists and selection boxes allow (i) the variable to be modelled; (ii) a single fixed effect; and (iii) one or more random effects to be chosen. A quantile-quantile plot assesses the fit of the variable (or its log-transformation) to the normal distribution. (b) The output of fitting a model containing the specified fixed and random effects is shown.

In the right-hand column of the tab users can select a single main factor or fixed effect to be tested for the selected variable. Below this, the **Select random factors** box allows multiple random factors to be added to the model; in mixed models these random factors can explain sources of variation which do not derive from the fixed effect. If no random factors are selected, the variable data will be fitted with a linear model using the lm function. If one or more random factors is selected, the data will be fitted with a linear mixed model using the lmer function. Note that if more than one random factor is selected, the factors will be nested in the order that they appear in the input CSV or Excel file.

At the bottom of the **Modelling** tab the output from the lm or lmer model fitting function is shown (Fig 4b). Estimates of statistical significance of the fixed effect are provided either by the output of lm or, in the case of mixed modelling, by the output of the Anova function from the car R package [13].

### Correlations

The top-level **Correlations** tab allows the investigation of associations between experimental variables (see Fig 5a). Pearson correlations are calculated between each pair of variables, calculated over all experimental units by default (but respecting the data selections made on the **Data Select** tab). Selecting an effect allows the user to subset the correlation calculation to those experimental units belonging to a particular level of a metadata factor (the effect and level are chosen from selection boxes in the sidebar at the left of the main plot). Correlations are drawn as a heat map, where colours run from blue (strong negative correlation) to red (strong positive correlation).

**Fig 5.**
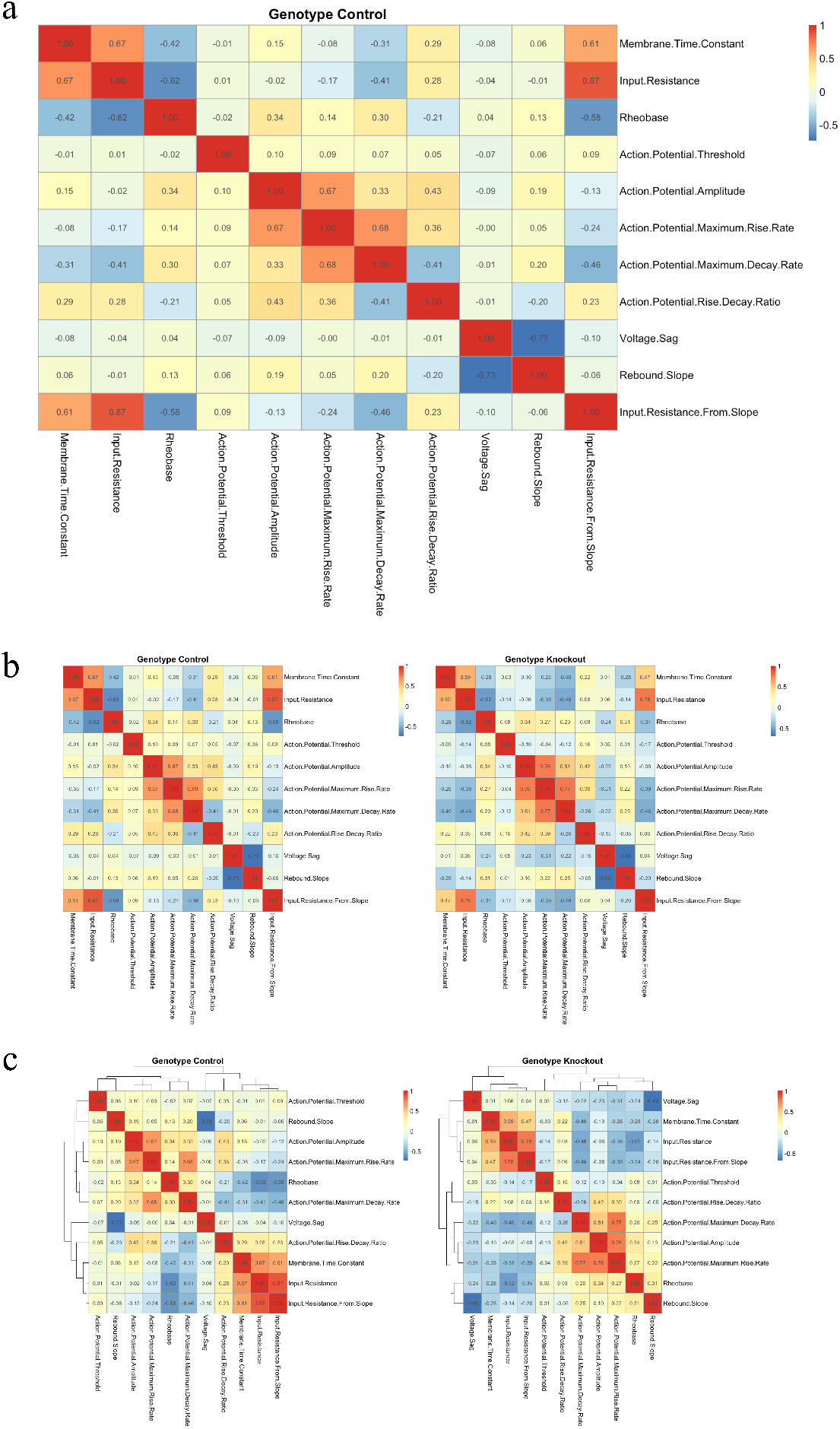
Heat maps in the Correlations tab show associations between experimental variables. (a) Here correlations between intrinsic electrophysiological properties are shown for neurons originating from control animals. For example, membrane time constant and input resistance are positively correlated, but voltage sag and rebound slope are strongly anti-correlated. (b) Paired correlation plots allow changes in variable associations to be examined. Here correlations between intrinsic electrophysiological properties are shown (i) for neurons originating from control animals, and (ii) for neurons originating from knockout mice. (c) Clustering groups strongly-related experimental variables. Correlations between intrinsic electrophysiological properties for control and knockout neurons are clustered, grouping together variables which show similar patterns of correlation.

If the **Pairwise** check box is selected then two heat maps are displayed, allowing the comparison of variable correlations calculated from different subsets of the data, defined by two levels of the chosen metadata factor (see Fig 5b). This allows changes in correlation between experimental conditions to be examined.

Finally, when the **Clustered** check box is selected then hierarchical clustering of experimental variables is performed, grouping together variables which show similar patterns of correlation against all the other variables (see Fig 5c). Otherwise, variables are displayed along the x- and y-axes of the heatmap in the order that they are defined in the input data file.

### Downloads

The top-level **Downloads** tab allows subsets of the input data to be exported as a new Excel spreadsheet or CSV file (see Fig 6a). The two boxes at the top of the page determine what combination of metadata and experimental variable columns will be present in the exported file; the table at the bottom of the page previews the data columns that will be exported. Data are exported via the **Download .xlsx** button as an Excel spreadsheet, or **Download .csv** button as a CSV file.

**Fig 6.**
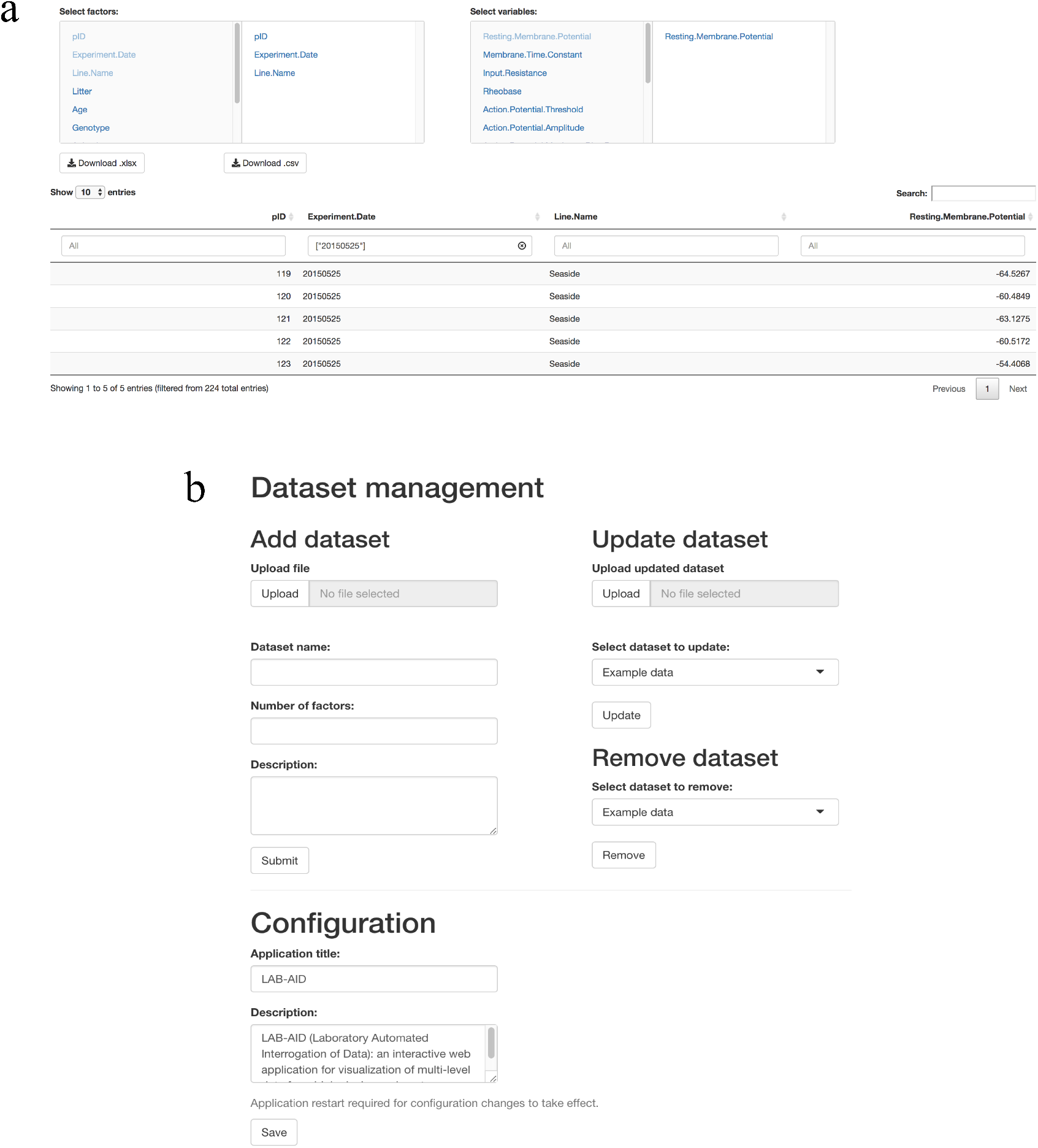
The Downloads and Configuration tabs allow for the export and import of data. (a) Subsets of the input data can be exported as Excel or CSV files. Here, three metadata columns and one experimental variable column have been selected for export. (b) The Configuration tab allows data sets to be added and removed. The application title and description can also be edited.

### Configuration

The top-level **Configuration** tab allows the user to add and remove data sets from within the application (see Fig 6b). To add a new data set, the user can upload a CSV file or an Excel spreadsheet containing data which adheres to the correct format (see **Methods**). Text fields allow the data set name, number of experimental factors, and data set description to be entered; while the data set description is optional, both data set name and the number of experimental factors must be provided for the data set to be successfully loaded. Newly added data can be immediately accessed on the **Data selection** tab. In the right column, the **Update dataset** interface allows users to upload an updated spreadsheet and select the name of the dataset to be replaced from a drop-down menu. To remove a data set, the user can simply select the data set from the **Remove dataset** drop-down menu and press the **Remove** button.

At the bottom of the tab there are options to change the application title, which is displayed at the left of the main tab bar, and the overall application description text, which appears on the top-level **About** tab. Note that changes to text on the **About** tab will take effect the next time the application is launched.

### Use cases

We briefly highlight, in the context of the demonstration electrophysiological data presented here, a number of situations that LAB-AID makes simple to examine:

1. After noting a neuron that is an outlier with respect to one electrophysiological measurement, one can choose that data point – by its unique pID – to be highlighted, and thus immediately see if the neuron is also an outlier with respect to other experimental variables.
2. By highlighting data points originating from particular animals or litters, for example (or those data points which are commonly identified by any other metadata factor), one can examine whether that animal or litter appears to be driving one or more apparent phenotypes.
3. By excluding data points originating from particular animals or litters (or, again, points which are commonly identified by any other metadata factor), one can examine whether correlations between electrophysiological measurements on neurons are robust to the removal of data originating from that animal or litter.

## Discussion

There is a great demand from biologists to be able to visualize the data from their laboratory experiments. Here we have presented LAB-AID, a simple, yet powerful, tool specifically designed to automatically visualize hierarchical data from biological experiments, that is freely available and easy to install and configure.

## Methods

### Technology

LAB-AID is implemented using Shiny [14], a package for the R programming language which allows interactive web applications to be easily built. Plots are produced with the ggplot2 [15], pheatmap [16] and patchwork [17] R packages, while interactive graph functionality is provided by the plotly package [18]. A full list of R packages required, and the versions that have been tested, is provided in Table 1.

**Table 1.** R packages used by LAB-AID.

| R package | Version tested | Reference |
| --- | --- | --- |
| car | 3.0-10 | [13] |
| jsonlite | 1.6 | [19] |
| lme4 | 1.1-23 | [12] |
| magrittr | 1.5 | [20] |
| patchwork | 0.0.1 | [17] |
| pheatmap | 1.0.10 | [16] |
| plotly | 4.8.0 | [18] |
| readxl | 1.1.0 | [21] |
| reshape2 | 1.4.3 | [22] |
| shiny | 1.1.0 | [14] |
| shinycssloaders | 1.0.0 | [23] |
| shinyjs | 1.0 | [24] |
| shinyWidgets | 0.4.5 | [25] |
| tidyverse (includes ggplot2) | 1.2.1 | [26] |
| WriteXLS | 4.0.0 | [27] |
| summarytools | 0.9.6 | [28] |

### Input data format

Data for LAB-AID should take the form of an Excel spreadsheet (both *.xls* and *.xlsx* files are supported) or a comma-separated values (CSV) file. Data tables should be in wide format, in which each row comprises metadata and measurements of experimental variables on a single experimental unit (for example, a cell). Columns containing metadata should appear first. In the parlance of mixed modelling, these will include both fixed and random effects; for example, our example data sheet contains a column indicating the genotype of the animal from which each measured neuron was derived (a fixed effect), and also columns indicating the brain slice and animal ID for each cell (random effects). Entries in metadata columns should be provided for every experimental unit.

Data columns should follow the metadata columns. Often, multiple data columns will be present, representing multiple simultaneous measurements made on the experimental unit. Not all measurements need to be recorded for every experimental unit.

### Installation and configuration

#### RShiny application installation

As LAB-AID is implemented using Shiny, it can be installed or deployed in the same way as any other Shiny application. For example, the code can be run on the user’s machine using RStudio [29], deployed to a web server using Shiny Server, or, if local server resources are not available, deployed to the cloud using shinyapps.io (https://www.shinyapps.io).

#### JSON configuration

LAB-AID configuration is stored in a simple JSON (JavaScript Object Notation, https://json.org) file residing in the same directory as the application code, which describes the location and structure of the input data. The configuration file describes a number of data sets, each of which corresponds to a particular Excel spreadsheet or CSV file. For each data set, the following must be defined:

1. **Name**: A name for the data set, which will appear in the LAB-AID user interface.
2. **Path**: Path to the Excel or CSV file containing data for this set, relative to the location of the LAB-AID code directory.
3. **n.Factors**: The number of metadata columns in this data set.
4. **Description** (optional): A description of this particular data set which will appear on the **About** tab.

Note that while all data sets defined in the configuration file will be available for exploration in the user interface, data sets do not need to contain the same metadata columns, nor do they need contain measurements on the same experimental units.

In addition, the main application title and description are also configured in the appropriate JSON fields:

1. **Title**: An overall title for the application which will appear at the left of the main tab bar.
2. **About**: An overall description of the data sets presented which will appear on the **About** tab.

The **Configuration** tab within the application provides an interface to add and remove data sets and to modify the application title and description. Alternatively, the configuration file can be edited manually.

## Availability

LAB-AID is freely available for download from GitHub under the MIT License, at https://github.com/sidbdri/LAB-AID.

## Author Contributions

Conceptualisation: P.K., O.D; Software Development: Z.K.; Experimental Data: S.B.; Writing, Original Draft: O.D., Z.K.; Writing, Review and Editing: Z.K., S.B., O.D., G.H., P.K.; Funding Acquisition: P.K.

